# Neutralization of Delta variant with sera of Covishield vaccinees and COVID-19 recovered vaccinated individuals

**DOI:** 10.1101/2021.07.01.450676

**Authors:** Gajanan N. Sapkal, Pragya D. Yadav, Rima R. Sahay, Gururaj Deshpande, Nivedita Gupta, Dimpal A Nyayanit, Deepak Y. Patil, Sanjay Kumar, Priya Abraham, Samiran Panda, Balram Bhargava

## Abstract

The recent emergence of B.1.617 lineage has created grave public health problem in India. The lineage further mutated to generate sub-lineages B.1.617.1 (Kappa), B.1.617.2 (Delta), B.1.617.3. Apparently, the Delta variant has slowly dominated the other variants including B.1.617.1 (Kappa), B.1.617.2 (Delta), B.1.617.3. With this, World Health Organization has described this sub-lineage as variant of concern. The high transmissibility associated with Delta variant has led to second wave of pandemic in India which affected millions of people. Besides this, variant of concerns has been reported to show lower neutralization to several approved vaccines. This has led to breakthrough infections after completion of vaccination regimen. There is limited information available on the duration of protective immune response post-infection, vaccination or breakthrough infection with SARS-CoV-2. In this study, we have evaluated immune response in sera of the Covishield vaccinated individuals belonging to category: I. one dose vaccinated, II. two doses vaccinated, III. COVID-19 recovered plus one dose vaccinated, IV. COVID-19 recovered plus two doses vaccinated and V. breakthrough COVID-19 cases. The findings of the study demonstrated that the breakthrough cases and the COVID-19 recovered individuals with one or two dose of vaccine had relatively higher protection against Delta variant in comparison to the participants who were administered either one or two doses of Covishield™. Prior vaccination results in less severe disease against subsequent infection provide evidence that both humoral and cellular immune response play an important role in protection.

---

The SARS-CoV-2 lineage B.1.617 was initially detected from India during October 2020 and since then further mutated as sub lineages B.1.617.1 (Kappa), B.1.617.2 (Delta) and B.1.617.3 variant. This has now mutated as Delta AY.1 and Delta AY.2, detected from India and many other countries. The Delta variant has been reported to be 60% more transmissible than the Alpha variant (B.111.7) and the WHO has designated Delta variant as a Variant of Concern (VOC). The second wave of the COVID-19 pandemic in India was dominated by Delta variant, affecting millions of people causing a serious public health crisis. Similarly, it spread rampantly and dominated over the Alpha variant in the UK as well and gained foothold in over 92 countries.^1^

The worldwide endeavor of scientists to create a safe and effective COVID-19 vaccine has resulted in the availability of 18 vaccines, which have received Emergency Use Authorization.^2^ The vaccines available against SARS-CoV-2, have shown efficacy ranging from 51 % to 94% against the original strain D614G in phase 3 clinical trials.^2^ Immune response to SARS-CoV-2 infection involves innate immune activation and antigen-specific responses of B and T cells.^3^ Particularly, the questions about immune escape of, newly emerging VOCs in vaccinated individuals are still being explored. For example, the efficacy of AZD1222, which was reported to be 70% in the UK and Brazil, only reached 22% in South Africa.^4^ Reduced efficacy against B.1.351 was also reported for the NVX-CoV237 (Novavax) and Ad26.COV2-S (Johnson & Johnson) vaccines.^4^

Currently, available vaccines appear to induce robust humoral and cellular immune responses against the SARS-CoV-2 spike protein.^4^ However, the newly emerged SARS-CoV-2 variants have led to breakthrough infections after completion of vaccination regimen.^5^ Hence, it is crucial to continuously evaluate vaccine-induced humoral immunity to SARS-CoV-2, immunity following natural infection, and the phenomenon of breakthrough infection to understand the immune escape due to emerging VOCs.

Covishield™ is a replication-deficient viral vector based-SARS-CoV-2 recombinant vaccine, which has been rolled out under the national COVID-19 vaccination program in India since 16 January 2021. Lacobucci et al., demonstrated significant immune responses following the first dose and complete seroconversion in the subjects after the second dose of Covishield™ in a study conducted in England and Wales.^6^ As the Delta variant has important mutations in spike region; it could pose a real challenge to the vaccines specifically developed targeting spike gene.

Earlier investigations from our group demonstrated a reduction in the neutralizing antibody (NAb) titer in the sera of Covishield™ vaccinees against B.1.617.1 (Kappa) variant.^7^ Here, we have assessed the NAb response of individuals immunized with Covishield™ (first dose and second dose), COVID-19 recovered individuals who were vaccinated (first dose and second dose) and breakthrough infections (due to Kappa and the Delta variant).

A comparative assessment of Covishield™ vaccinated individuals’ (n=116) sera in different categories was performed against prototype strain B.1 (D614G) and Delta variant. Sera under this study were grouped into five categories: I. one dose (n=31), II. Two doses (n=31), III. COVID-19 recovered plus one dose (n=15), IV. COVID-19 recovered plus two doses (n=19) and V. breakthrough COVID-19 cases (n=20). All the sera were collected four weeks post-vaccination for category I-IV participants. For category V patients, sera were collected two weeks after completing a two-dose immunization schedule and were SARS-CoV-2 positive through RT-PCR. COVID-19 recovered cases (category III & IV) were of B.1 lineage whereas breakthrough cases (category V) belonged to lineages Kappa and Delta variants which were confirmed by next-generation sequencing. The samples were also screened for receptor binding domain-spike RBD-S1 and N protein ELISA (Supplemental information). Except for category I and II samples were not screened for N protein ELISA.

The NAb titers against B.1 and B.1.617.2 strains were determined for sera of each category using plaque reduction neutralization test (Supplemental information). NAb against B.1 were not observed in 11/31 (35.5%) participants in category I. Similarly, NAb against the B.1.617.2 were not observed in 18/31 (58.1%) and 5/31 (16.1%) participants of categories I and II respectively. The GMT ratio of the B.1.617.2 vs B.1 for categories I- V were 0.22 (95% confidence interval (CI): 0.19-0.25; p-value <0.0001); 0.31 (95% CI: 0.22-0.43; p-value <0.0001); 0.34 (95% CI: 0.32-0.37; p-value <0.0001); 0.38 (95% CI: 0.38-0.39; p-value <0.0001), and 0.53 (95% CI: 0.49-0.56; p-value <0.0001) respectively (Figure 1 A-E). Wilcoxon signed rank matched-pairs test was used to assess the statistical significance of the difference in NAb titers.

**Figure 1.**
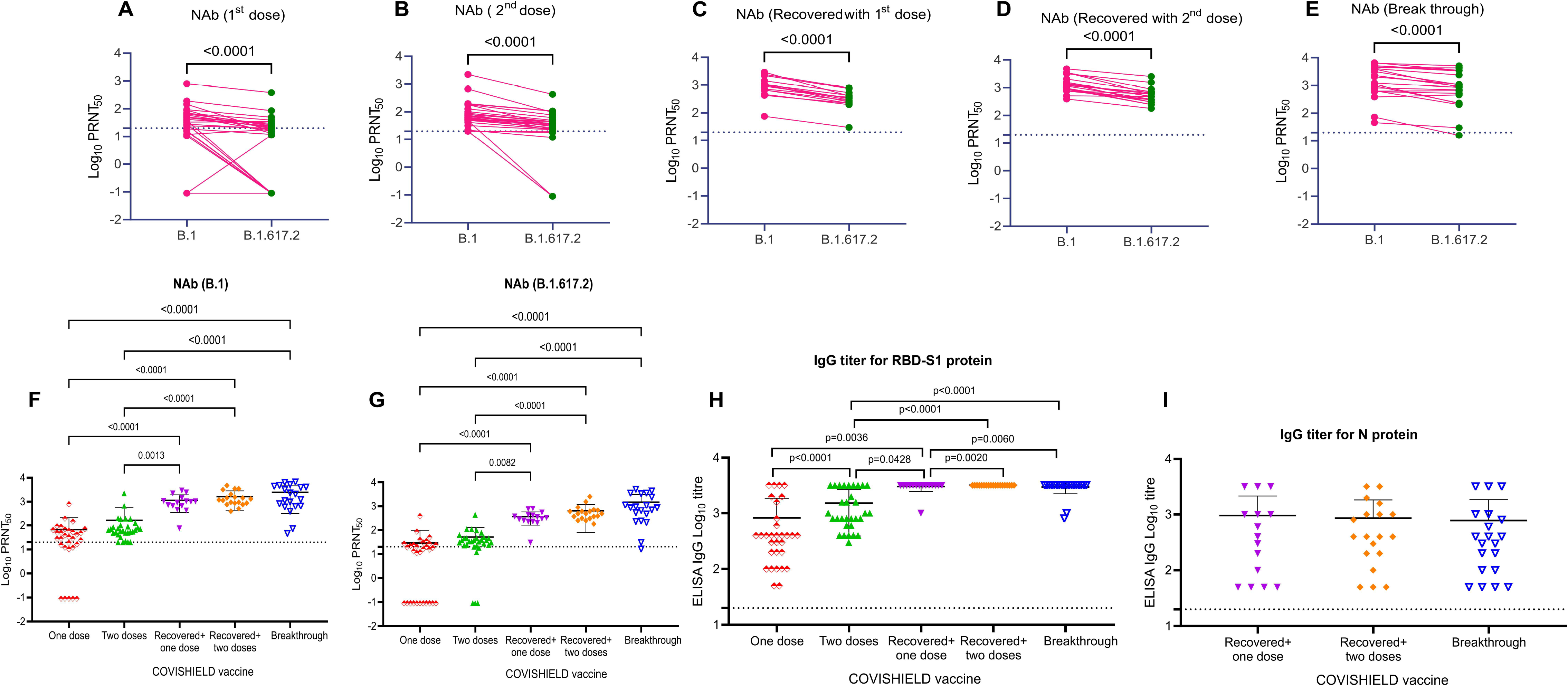
Neutralization antibody titer of individual sera from different category against SARS-CoV-2 B.1 and B.1.617.2 strains and RBD-S1 and N protein titer. NAb titer against the B.1 (pink) and B.1.617.2 (green) was compared between participants administered with one dose Covishield™ vaccine (A), two doses vaccine (B) COVID-19 recovered individuals administered with one dose Covishield™ vaccine (C) COVID-19 recovered individuals administered with two doses (D) and breakthrough participant (E). A two-tailed pair-wise comparison was performed using the Wilcoxon matched-pairs signed-rank test to analyze the statistical significance. Comparison of NAb titer between participants administered with one dose Covishield™ vaccine (red color), two doses Covishield™ vaccine (green color) Covishield™ vaccine along COVID-19 recovered individuals administered with one dose (purple color) and two doses (indigo color) and breakthrough (black color) participants against B.1 strain (F) and B.1.617.2 strain (G). IgG titers of participant’s sera from different categories for SARS-CoV-2 RBD protein (H) and N protein ELISA (I). The statistical significance was assessed using two-tailed Kruskal Wallis test with Dunn’s test of multiple comparisons to analyze the statistical significance and a p-value less than 0.05 was considered to be statistically significant. The dotted line on the figures indicates the limit of detection of the assay. Data are presented as mean values +/− standard deviation (SD).

NAb titers for B.1.617.2 relative to B.1 were reduced in the sera of the participants belonging to categories I (78%), II (69%), III (66%), IV (38%) and V (47%). This reduction in NAb titers in the sera of participants lends support to increased susceptibility of the population who has been immunized with even both doses of the vaccine with Delta variant.

The GMT for B.1 strain in categories I-V was 16.12 (95%CI: 6.507-39.94; p-value <0.0001), 73.47 (95% CI: 50.7-106.5; p-value <0.0001), 868.9 (95% CI: 533.3-1416; p-value >0.999), 1312 (95%CI: 949.3-1813; p-value >0.999) and 1344 (95%CI: 700.2-2580) respectively. This indicates that sera of COVID-19 recovered participants who were vaccinated with either one (category III) or two doses (category IV) and breakthrough had higher NAb titers relative to COVID-19 negative participants of category I and II (Figure 1F). The significantly higher NAb titers in sera of participants of COVID-19 recovered (categories III and IV) as compared to COVID-19 negative (categories I and II) highlights the fact that even one dose of vaccine in convalescent patients is enough to provide effective protection against re-infection of SARS-CoV-2 or protection against newly emerging variants. Similar conclusions have been demonstrated by studies supporting single-dose vaccination after recovery from previous infection.^8^ This finding supports the role of cross-reactive SARS-CoV-2-specific T-cell-mediated immune response as has been described by Geers et. al.^9^

Similarly, the GMT for B.1.617.2 strain in categories I-V was 3.553 (95%CI: 1.252-10.08), 22.43 (95%CI: 10.96-45.9), 298.8 (95%CI: 196-455.5), 501.3 (95%CI: 368.6-681.7) and 706.2 (95%CI: 342.8-1455) respectively (Figure 1G). Participants in category I (4.5 fold) and II (3.2 fold) showed reductions in NAb titers against Delta variants as compared to B.1 lineage. Reduction in GMT was evident in categories III-V, however the NAb titers remained significantly higher to provide enough protection. An increase in the NAb titers was observed for both B.1 and B.1.617.2 strains in the sera of the participants who had completed two vaccine doses relative to one dose. NAb titers against B.1 and B.1.617.2 strains were highest among breakthrough participants which may be due to spike-specific T-cell responses.^10^ A two-tailed Kruskal Wallis test with Dunn’s test of multiple comparisons was used to analyze the statistical significance.

IgG titer specific to SARS-CoV-2 RBD and N-protein was analyzed for participants in different categories. It was observed that IgG specific to RBD protein showed higher antibody response (1:3200) in category III-V group (Figure 1 H). N protein-based ELISA indicated a similar pattern of IgG titer in participants in category III-V (Figure 1 I).

We observed significantly lower NAb titers for the B.1.617.2 strain relative to B.1 strain. However, NAbs in breakthrough participants and the COVID-19 recovered individuals with one or two-dose of vaccine had relatively higher protection against B.1.617.2 in comparison to the participants who were administered either one or two doses of Covishield™. In addition to direct virus neutralization, antibodies can have multiple other modes of action that are primarily mediated by IgG1 and IgG3 subclass antibodies.^11^ Priming of the immune system memory is established by natural infection or vaccination. Spike-specific IgG antibodies and virus-specific memory T cells may decrease with time, but virus-specific memory B-cells increase.^12^

Long-term follow-up of participants could help understand the impact of natural infection and vaccination on long-term protection from SARS-CoV-2 offered by Covishield™. It is important to track the breakthrough infections to look for unexpected changes. The limitation of this study includes the unavailability of data on cell-mediated responses which may be important for cross-reactive protection. The third dose of ChAdOx1 nCoV-19 had significantly boosted the antibody titers and Spike-specific T-cell responses above the second dose.^10^

Monitoring of breakthrough infection would make us understand the impact of new variant or VOC on escape of vaccine induced immunity. Data has shown again and again that if the individuals get infected post vaccination, had been protected from severe disease.

## Supporting information

Supplementary Information Methodology

## Ethical approval

The study was approved by the Institutional Human Ethics Committee of ICMR-NIV, Pune, India under project ‘Assessment of immunological responses in breakthrough cases of SARS-CoV-2 in post COVID-19 vaccinated group’.

## Author Contributions

GNS, PDY and PA contributed to study design, data analysis, interpretation and writing and critical review. RRS, GRD, DAN, DYP and SK contributed to data collection, data analysis, interpretation, writing and critical review. NG, SP, and BB contributed to the critical review and finalization of the paper.

## Acknowledgement

We Sincere acknowledge the excellent support of Dr. Rajlaxmi Jain, Mr. Prasad Sarkale, Mr. Shreekant Baradkar, Ms. Aasha Salunkhe, Mr. Chetan Patil, Mrs. Triparna Majumdar and Mrs. Savita Patil for extending excellent technical support.

